# Proteoform identification using multiplexed top-down mass spectra

**DOI:** 10.1101/2025.02.05.636727

**Authors:** Zhige Wang, Xingzhao Xiong, Xiaowen Liu

**Affiliations:** Department of Computer Science, Tulane University, New Orleans, Louisiana, 70112, United States; Deming Department of Medicine, Tulane University, New Orleans, Louisiana, 70112, United States

## Abstract

Top-down mass spectrometry (TDMS) is the method of choice for analyzing intact proteoforms, as well as their post-translational modifications and sequence variations. In TDMS experiments, multiple proteoforms are often co-fragmented in tandem mass spectrometry (MS/MS) analysis, resulting in multiplexed TD-MS/MS spectra. Since multiplexed TD-MS/MS spectra are more complex than common spectra generated from single proteoforms, these spectra pose a significant challenge for proteoform identification and quantification. Here we present TopMPI, a new computational tool specifically designed for the identification of multiplexed TD-MS/MS spectra. Experimental results demonstrate that TopMPI significantly increases proteoform identifications and reduces identification errors in multiplexed TD-MS/MS spectral analysis compared to existing tools.

## Introduction

Top-down mass spectrometry (TDMS) has gained increasing attention in proteomics due to its ability to directly examine intact proteoforms and characterize proteoforms with various alterations arising from gene mutations, alternative splicing, and post-translational modifications (PTMs) [1, 2]. Unlike bottom-up MS, in which combinatorial PTM patterns on proteoforms are lost during enzymatic digestion prior to MS, TDMS enables the analysis of these patterns along with other sequence variations, facilitating the study of their biological functions [3].

In a typical TDMS experiment, MS1 spectra are generated to profile proteoforms in the sample, and data-dependent acquisition (DDA) [3] or data-independent acquisition (DIA) [4] is employed to isolate and fragment specific proteoform ions within predefined isolation windows to generate tandem mass spectrometry (MS/MS) spectra. Multiplexed MS/MS spectra containing fragment ions from co-fragmented proteoforms are frequently observed [5], especially in top-down DIA-MS [4], adding a layer of complexity to TDMS data analysis.

Many methods have been proposed for the identification of multiplexed DIA mass spectra in bottom-up proteomics [6-9]. When spectral libraries are available, spectra from the library are matched to bottom-up multiplexed mass spectra for peptide identification [7, 10]. Otherwise, multiplexed mass spectra are demultiplexed based on the similarity of retention time profiles of precursor and fragment ions [6]. Subsequently, the demultiplexed spectra are searched against a protein database for peptide identification.

Both database searching and spectral library searching have been used to identify multiplexed DDA mass spectra in bottom-up proteomics [8, 9, 11]. One main difference between DIA and DDA multiplexed mass spectra is that the demultiplexing approach used in DIA-MS is ineffective for DDA-MS because fragment ion retention time profiles are unavailable for most multiplexed DDA spectra.

Several top-down DIA-MS studies were reported recently [4, 12] and the demultiplexing method has been extended to analyze top-down multiplexed DIA-MS data [4]. However, due to limitations in top-down DIA-MS data analysis, DDA-MS remains the predominant method in top-down proteomics [2].

There remains a lack of software tools for identifying top-down multiplexed DDA mass spectra. Many computational tools, like MSPathFinder [13], ProSightPD [14],TopPIC [15], and TopMG [16], have been developed for top-down spectral identification through database searching. While these tools are efficient in identifying MS/MS spectra generated from single proteoforms, they are unable to identify multiple proteoforms from multiplexed spectra. Spectral library-based methods used in bottom-up MS can be adapted for top-down MS [9], but they rely on comprehensive spectral libraries. Additionally, database search methods [8, 9, 11] for analyzing multiplexed bottom-up DDA-MS data are inefficient for processing multiplexed top-down DDA-MS data due to the complexity of top-down mass spectra.

To address this challenge, we introduce TopMPI (TOP-down mass spectrometry-based Multiple Proteoform Identification), a new software tool designed to identify multiple proteoforms from multiplexed TD-MS/MS spectra via database searching. Experimental results demonstrate that TopMPI significantly improves proteoform identification from top-down DDA-MS data of complex biological samples compared to existing software tools, which focus solely on identifying single proteoforms from top-down DDA mass spectra.

## Methods

### *E. coli* sample preparation

*E. coli* K12 MG1655 cells were pelleted by centrifugation at 5,000×g and 4°C for 5 min and then washed with 5mL 1x PBS. The cell pellets were resuspended in 200 μL of 25 mM ammonium bicarbonate (ABC) buffer with the addition of 1× (v/v) EDTA-free protease inhibitor. Then the cell pellets were lysed using 0.1 mm beads, which were mixed with the cell suspension and ABC buffer in a 1:1:2 (v/v) ratio, followed by bead beating for 3 min. After bead beating, the cell lysate was centrifuged at 12,000×g and 4°C for 4 min to remove insoluble debris. Proteins were filtered using Amicon Ultra-0.5 filters by centrifugation at 14,000×g and 4°C for 20 min, retaining only proteins with a molecular weight between 3 kDa and 100 kDa. Next, 1 μL of 1M dithiothreitol (DTT) was added to the lysate and incubated at 55°C for 45 min. Then, 2.5 μL of 1M iodoacetamide (IAA) was added and allowed to react at room temperature for 30 min. The protein concentration was determined using the Pierce BCA Protein Assay Kit.

### Top-down RPLC-MS/MS analysis

A total of 300 ng of *E. coli* protein was analyzed using a Thermo Scientific (Waltham, MA, USA) Ultimate 3000 reverse-phase liquid chromatography (RPLC) system equipped with a C2 column (100 μm i.d., 60 cm length, CoAnn, Richland, WA, USA) and coupled to a Thermo Orbitrap Lumos mass spectrometer (Waltham, MA, USA). Mobile phase A was water with 0.1% formic acid (FA). Mobile phase B was 60% acetonitrile (ACN), 15% isopropanol (IPA) and 25% water with 0.1% FA. A 98-min gradient (0-5 min 5%, 5-7 min for 5% to 35%, 7-10 min for 35% to 50%, 10-97 min for 50% to 80%, 97-98 min for 80% to 99%) was applied for proteoform separation with a flow rate of 400 nL/min.

MS1 scans were collected at a resolution of 240,000 (at 200 *m/z*) with 4 microscans, a maximum injection time of 200 ms, and a scan range of 720-1200 *m*/*z*. The top 6 precursors in each MS1 scan were selected for Higher-energy C-trap dissociation (HCD) MS/MS analyses with following settings: the precursor isolation window was 3 *m*/*z*, the normalized collision energy was 30%; the Automated Gain Control (AGC) target was 10^6^, and the maximum injection time was 500 ms; the microscan number was 1; the resolution was 60,000 (at 200 *m/z*); and the scan range was 400-2000 *m*/*z*.

### Top-down mass spectral preprocessing

Top-down MS raw data files were converted to mzML files using msconvert [17], and then the mzML files were analyzed by TopFD [18] (version 1.7.6 and Supplemental Table S1 for parameter settings) for spectral deconvolution and feature detection. For each reported proteoform feature with multiple charge states, a single charge proteoform feature (SCPF) was generated for each charge state. For each MS/MS spectrum, the SCPFs detected within its isolation window were ranked according to their total peak intensities within the window, and the two most intense SCPFs were assigned to the spectrum.

### Primary precursor selection

An MS/MS spectrum is classified as a multiplexed spectrum if the ratio between the total peak intensities of the second and first most abundant SCPFs in the isolation window exceeds *α*, where *α* is a user-specified parameter with a default setting of 20%. For a multiplexed spectrum containing two precursors, two rounds of database searches are conducted to identify two proteoforms. The precursor used in the first round of the database search is referred to as the primary precursor, while the other is designated as the secondary precursor. Notably, the primary precursor may have a lower intensity than the secondary precursor.

For a deconvoluted MS/MS spectrum *S* with two SCPFs *F*_1_ and *F*_2_, where *F*_1_ has a higher total peak intensity in the isolation window than *F*_2_, the primary precursor is determined as follows.

Let *S*_1_ be the spectrum with precursor *F*_1_ and all fragment masses in *S*, and let *S*_2_ be the spectrum with precursor *F*_2_ and all fragment masses in *S*. The two spectra are searched against the corresponding protein sequence database for proteoform identification using TopPIC (version 1.7.6) without filtering of reported PrSMs (Fig. 1a). If TopPIC reports only one PrSM from the two spectra, then precursor of the spectrum of the PrSM is the primary one. If a PrSM is reported for each of two spectra, we will first check if the two PrSMs are consistent and then determine the primary precursor.

**Fig. 1:**
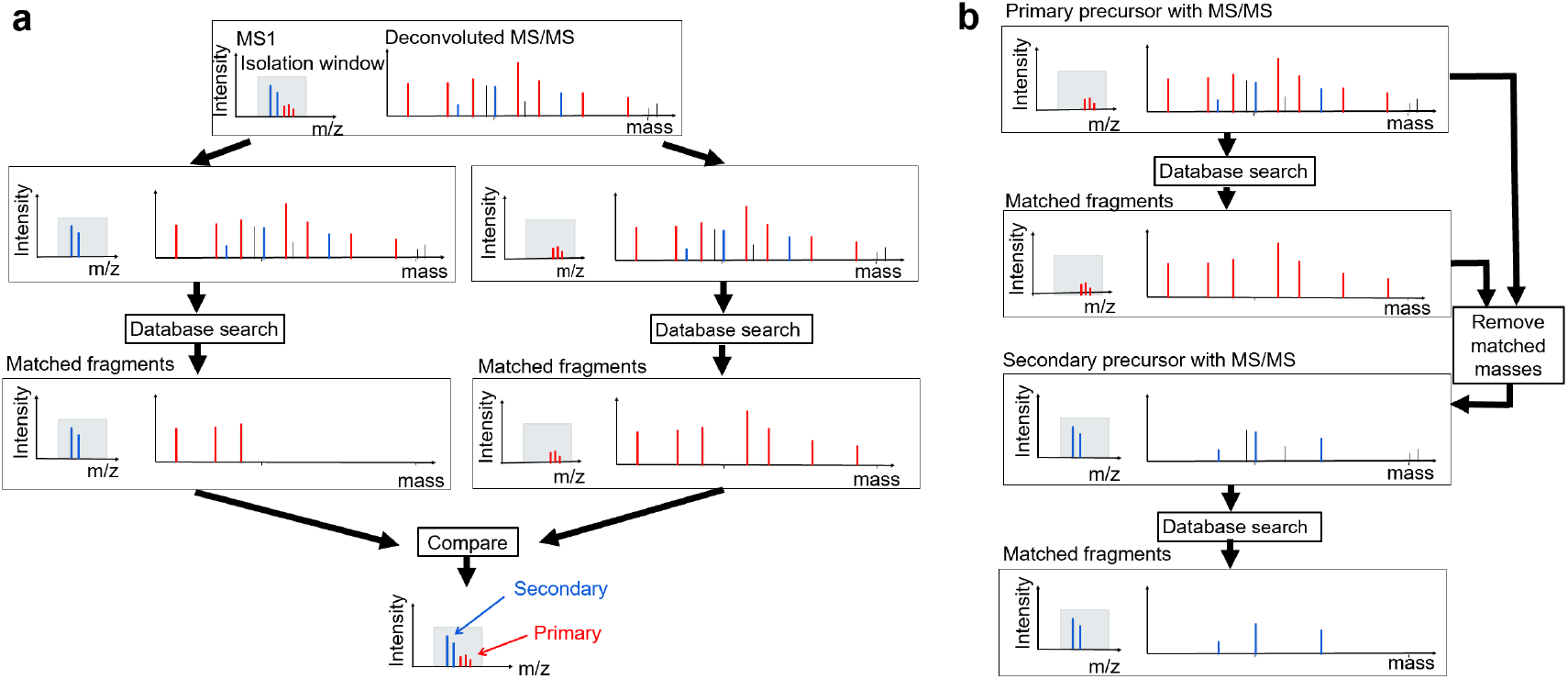
The overview of TopMPI. (a) Primary precursor selection for a multiplexed MS/MS spectrum with two precursors (blue and red) in the MS1 spectrum, along with their corresponding fragment masses (blue and red) and noise fragments (black). The red and blue precursors, along with their respective fragment masses, are independently searched against a protein sequence database, resulting in a correct identification for the red precursor and an incorrect identification for the blue precursor. The red precursor is selected as the primary precursor because its identification has a higher number of matched fragment masses. (b) Two-round database search strategy. The primary precursor (red) and all fragment masses in the spectrum are initially searched against the protein sequence database for proteoform identification. Once a match is found, the matched fragment masses (red) are removed. The secondary precursor (blue) and the remaining fragment masses are then searched against the protein sequence database for proteoform identification.

Two PrSMs are inconsistent if they are matched to the same protein or most of their matched fragment masses are shared. Let *M*_1_ and *M*_2_ represent the sets of matched experimental fragment masses in the two PrSMs reported by the database search, respectively. The two PrSMs are inconsistent if 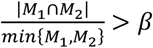, where *β* is a user specified parameter (*β* = 0.9 in the experiments).

Inconsistency between two PrSMs can arise in two scenarios. First, the two features *F*_1_ and *F*_2_ are from two similar proteoforms of the same protein, with most fragment masses overlapping. Second, the two features *F*_1_ and *F*_2_ are from two proteins, and one of the two PrSMs is incorrect. If the PrSMs of *S*_1_ and *S*_2_ are consistent, then *F*_1_ is chosen as the primary precursor.

Otherwise, the primary precursor is determined using the following method. We first normalize the number of matched fragment masses for each PrSM based on the presence of unknown mass shifts. For a PrSM with *x* matched fragment masses and *y* unknown mass shifts, the number of normalized matched fragment masses (NMFMs) is computed as *x*-*yδ*, where *δ* is a user specified parameter (*δ*=5 in the experiments). If the PrSM of *S*_2_ has at least *γ* more NMFMs than that of *S*_1_, where *γ* is a user specified parameter (*γ* = 4 in the experiments), then *F*_2_ is chosen as the primary precursor. Otherwise *F*_1_ is the primary precursor.

### Proteoform identification by multiplexed top-down DDA mass spectra

TopMPI employs a method similar to CharmeST [11] to identify two proteoforms from multiplexed top-down DDA mass spectra (Fig. 1b). For an MS/MS spectrum *S* with a primary precursor *F*_1_ and a secondary precursor *F*_2_, a two-round search is conducted to identify two proteoforms. In the first round, the primary precursor *F*_1_ and the fragment masses in spectrum *S* are searched against the corresponding protein sequence database, concatenated with a decoy database, using TopPIC (version 1.7.6). If a PrSM is reported, the matched fragment masses of the PrSM are removed from *S*. In the second round, the secondary precursor and the remaining fragment masses in *S* are searched against the same target-decoy protein sequence database using TopPIC. The PrSMs identified in the first and second rounds are filtered separately using a 1% spectrum-level false discovery rate (FDR). The filtered PrSMs from both rounds are then merged, and the identified PrSMs are clustered into proteoform groups. Two PrSMs are assigned to the same group if their precursors are from the same proteoform feature reported by TopFD or if they are matched to the same protein with a precursor mass difference of less than 1.2 Da. Finally, the identified proteoforms are filtered using a 1% proteoform-level FDR.

## Results

### Overview of TopMPI

TopMPI is designed to identify two proteoforms from a multiplexed top-down DDA MS/MS spectrum with two precursors. Its primary function is to determine the order of the two precursors for database search-based proteoform identification (Fig. 1a). A common error in multiplexed top-down mass spectral identification occurs when the most abundant precursor is incorrectly combined with fragment masses from a different precursor, leading to incorrect proteoform identification. TopMPI mitigates such errors by swapping the order of the two precursors before performing the database search (see Methods).

The precursor selected for the first round of the database search is designated as the primary precursor, while the other is assigned as the secondary precursor. Once the primary and secondary precursors are determined, TopMPI employs a method similar to CharmeST [11] to identify two proteoforms from a multiplexed top-down DDA mass spectrum (Fig. 1b). TopMPI first uses the primary precursor and all the fragment masses in the MS/MS spectrum to identify a PrSM by database search. Following identification, the matched fragment masses in the reported PrSM are removed from the spectrum, and a new spectrum is generated using the secondary precursor and the remaining fragment masses for spectral identification (see Methods).

### Precursor intensity and spectral identification

A top-down DDA-MS dataset containing 10,320 MS/MS spectra was generated from *E. coli* K12 MG1655 cells (see Methods) to investigate how precursor intensities in the isolation window influence the identification of multiplexed spectra. Following data preprocessing (see Methods), TopPIC (version 1.7.6; parameter settings provided in Supplemental Table S2) was used to search the MS/MS spectra against the UniProt *E. coli* K12 proteome database (version September 7, 2023; 4,530 entries) concatenated with a decoy database of equal size. In TopPIC, for each MS/MS spectrum, only the most abundant precursor within the isolation window was combined with fragment masses for proteoform identification. Once the first PrSM was reported, the matched fragment masses corresponding to the identified proteoform were removed, and a new MS/MS spectrum was generated using the second most abundant precursor and the remaining fragment masses. This newly generated spectrum was then searched against the same target-decoy database using TopPIC with identical parameter settings. The PrSMs reported from the most abundant and second most abundant precursors were filtered separately using a 1% spectrum-level FDR.

A total of 1,991 and 170 PrSMs were identified from the most and the second most abundant precursors, respectively, with 86 spectra containing two reported PrSMs. The intensity ratio between the second and first most abundant precursors was calculated for each of these 86 spectra (Supplemental Fig. S1), and the distribution indicated that all ratios exceeded 20%. Based on this observation, the parameter *α* for detecting multiplexed spectra was set to 0.2 (see Methods). Consequently, in these experiments, a spectrum was classified as multiplexed only if the intensity ratio between the second and first most abundant precursors was at least 20%.

### Evaluation on pseudo-multiplexed MS/MS spectra

To assess the performance of TopMPI in identifying multiplexed spectra, evaluation datasets of pseudo-multiplexed MS/MS spectra were generated from *the E. coli* dataset. TopPIC (version 1.7.6) was used to search the MS/MS spectra against the UniProt *E. coli* K12 proteome database (version September 7, 2023; 4,530 entries) and identified 1,779 PrSMs using an E-value cutoff of 0.01 (parameter settings detailed in Supplemental Table S3). Since some of the identified MS/MS spectra were multiplexed, the spectra were filtered based on the SCPFs observed in their isolation windows in MS1 spectra. For each of the 1,779 MS/MS spectra, the SCPFs within its isolation window were ranked according to the sum of their peak intensities. If the intensity sum of the most abundant SCPF was less than 85% of the total peak intensity sum of all SCPFs, the MS/MS spectrum was excluded, yielding 498 spectra, which were treated as non-multiplexed spectra. A non-multiplexed spectrum was classified as a zero-shift spectrum if its proteoform contained no unknown mass shifts and as a one-shift spectrum if its proteoform contained one unknown mass shift. Among the 498 spectra, 251 were zero-shift spectra, and 247 were one-shift spectra.

To generate a pseudo-multiplexed MS/MS spectrum, a non-multiplexed experimental spectrum was first selected as the base spectrum, and another non-multiplexed spectrum was chosen as the additive spectrum. Some fragment masses from the additive spectrum were then added to the base spectrum. A spectrum was considered a valid additive spectrum for a given base spectrum if it satisfied three criteria: (1) the distance between the SCPF average *m/z* values of the additive and base spectra was no greater than 20 *m/z*, (2) the proteoform identifications of the base and additive spectra are from two different proteins, and (3) after removing from the additive spectrum the fragment masses that are randomly matched to the b- or y-ions of the proteoform of the base spectrum, the number of remaining fragment masses in the additive spectrum was at least twice that of the base spectrum.

Among the 498 spectra, 256 had both a valid zero-shift additive spectrum and a valid one-shift additive spectrum. Of these 256 base spectra, 134 were zero-shift spectra, and 122 were one-shift spectra. The 256 base spectra and their corresponding zero-shift additive spectra were used to generate 256 spectral pairs. The spectral pairs in which the base spectrum was a zero-shift or one-shift spectrum were referred to as Base0-Add0 and Base1-Add0 pairs, respectively. Similarly, the 256 base spectra and their corresponding one-shift additive spectra were used to generate an additional 256 spectral pairs, and those with zero-shift and one-shift base spectra were referred to as Base0-Add1 and Base1-Add1 pairs, respectively. In total, 512 spectral pairs were generated. Within each pair, the base spectrum precursor was treated as the most abundant precursor in the multiplexed spectrum, and the protein identifications of the base and additive spectra were referred to as the base protein and additive protein, respectively.

The 512 spectral pairs were used to generate 11 pseudo-multiplexed spectrum data sets, each corresponding to an additive-base ratio (A-B ratio) of *r* = 0%, 20%, 40%, …, 200%. The data set with an A-B ratio of *r* contained 512 spectra, where each spectrum included all fragment masses from the base spectrum and a randomly selected subset of round(*rn*) fragment masses from the additive spectrum, where *n* represents the number of fragment masses in the base spectrum, and round() denotes the rounding function.

We evaluated the error rates of TopMPI under different parameter settings for *δ* and *γ* using the pseudo-multiplexed spectrum dataset with *r* = 100%. The mass spectra were searched against the UniProt *E. coli* K12 proteome database (version September 7, 2023; 4,530 entries) concatenated with a shuffled decoy database of the same size using TopMPI (parameter settings provided in Supplemental Table S4). The identified PrSMs were filtered using a 1% spectrum-level FDR. Two types of incorrect spectral identifications were reported by TopMPI. The first type, precursor selection errors (PSEs), occurred when the precursor of the base spectrum was incorrectly assigned to a proteoform of the additive protein, or *vice versa*. The second type, random matching errors (RMEs), occurred when the precursor of the base or additive spectrum was matched to a protein that was neither the base protein nor the additive protein. A comparison of TopMPI’s performance under various *δ* and *γ* parameter settings showed that the lowest PSE rate was achieved when *δ* = 5 and *γ* = 4 (Fig. 2), which were selected as the default settings for TopMPI. Additionally, the settings of *δ* and *γ* did not significantly impact the RME rates.

**Fig. 2:**
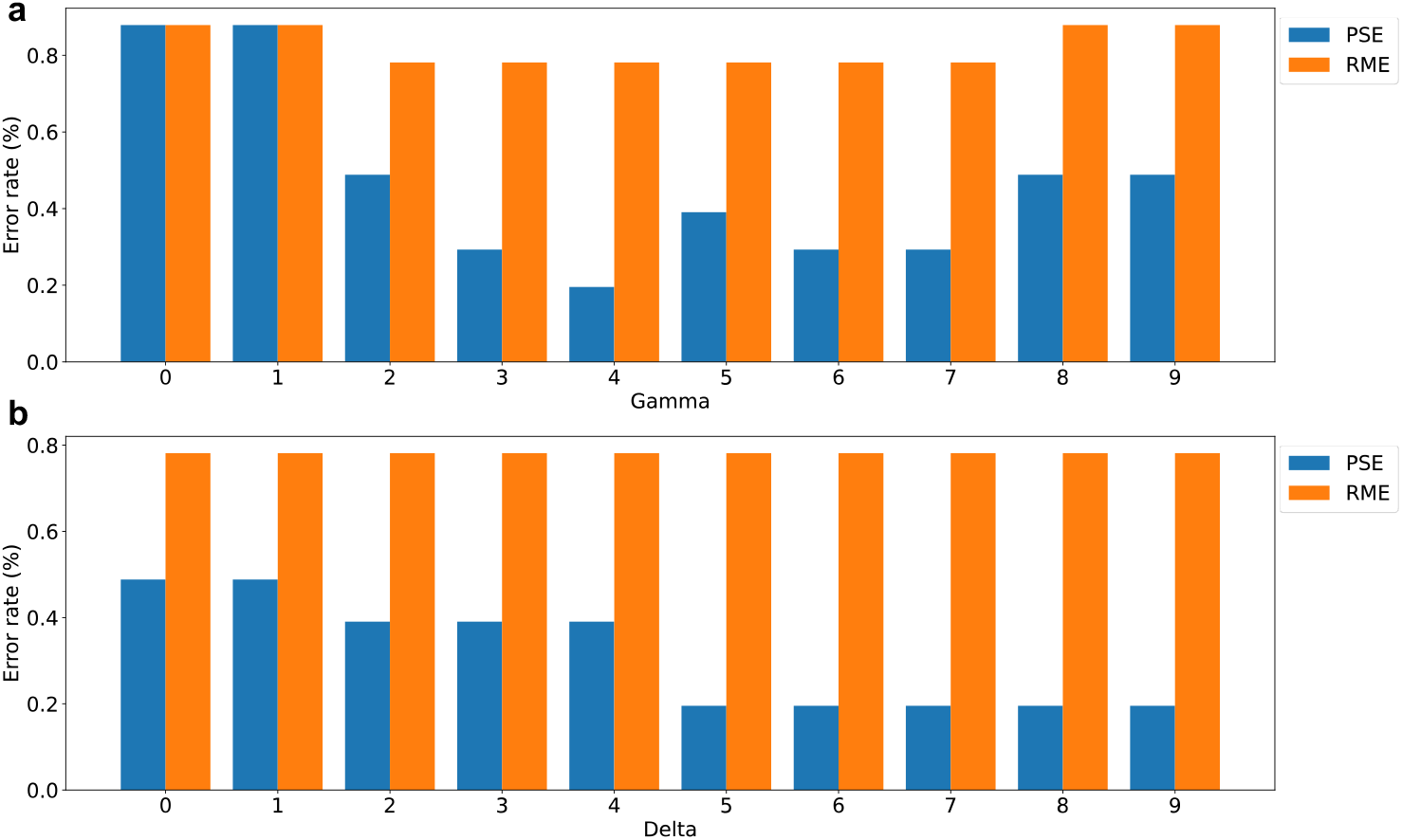
PSE and RME rates of TopMPI with various *δ* and *γ* parameter settings on the pseudo-multiplexed spectrum dataset generated with *r* = 100%. (a) Error rates with *δ* = 5 and varying *γ* values (0, 1, …, 9). (b) Error rates with *γ* = 4 and varying *δ* values (0, 1, …, 9).

To benchmark the accuracy of spectral identifications, we compared TopPIC (version 1.7.6) and TopMPI using the 11 datasets. The mass spectra were searched separately against the target-decoy UniProt *E. coli* K12 proteome database using TopPIC and TopMPI (parameter settings provided in Supplemental Tables S2 and S5). The identified PrSMs were filtered using a 1% spectrum-level FDR. As the A-B ratio increased, the error rates of spectral identifications reported by TopPIC also increased across the four types of spectrum pairs (Fig. 3a and 3b). The majority of errors reported by TopPIC were PSEs, with higher error rates when the proteoform of the base spectrum contained unknown mass shifts. For base precursors, TopMPI significantly reduced PSEs in spectral identifications (Fig. 3a) while reporting a similar number of RMEs (Fig. 3b) compared to TopPIC. However, the identification sensitivity of TopMPI was slightly lower than that of TopPIC (Fig. 3c). One possible explanation for this reduction in sensitivity is that TopMPI removes some identifications to correct PSEs. When a base precursor is incorrectly matched to the additive protein, TopMPI swaps the order of the base and additive precursors in the database search to correct the error, thereby also removing the incorrect identification of the base precursor. Furthermore, TopMPI exhibited low error rates for spectral identifications from the additive precursors (Fig. 3d).

**Fig. 3:**
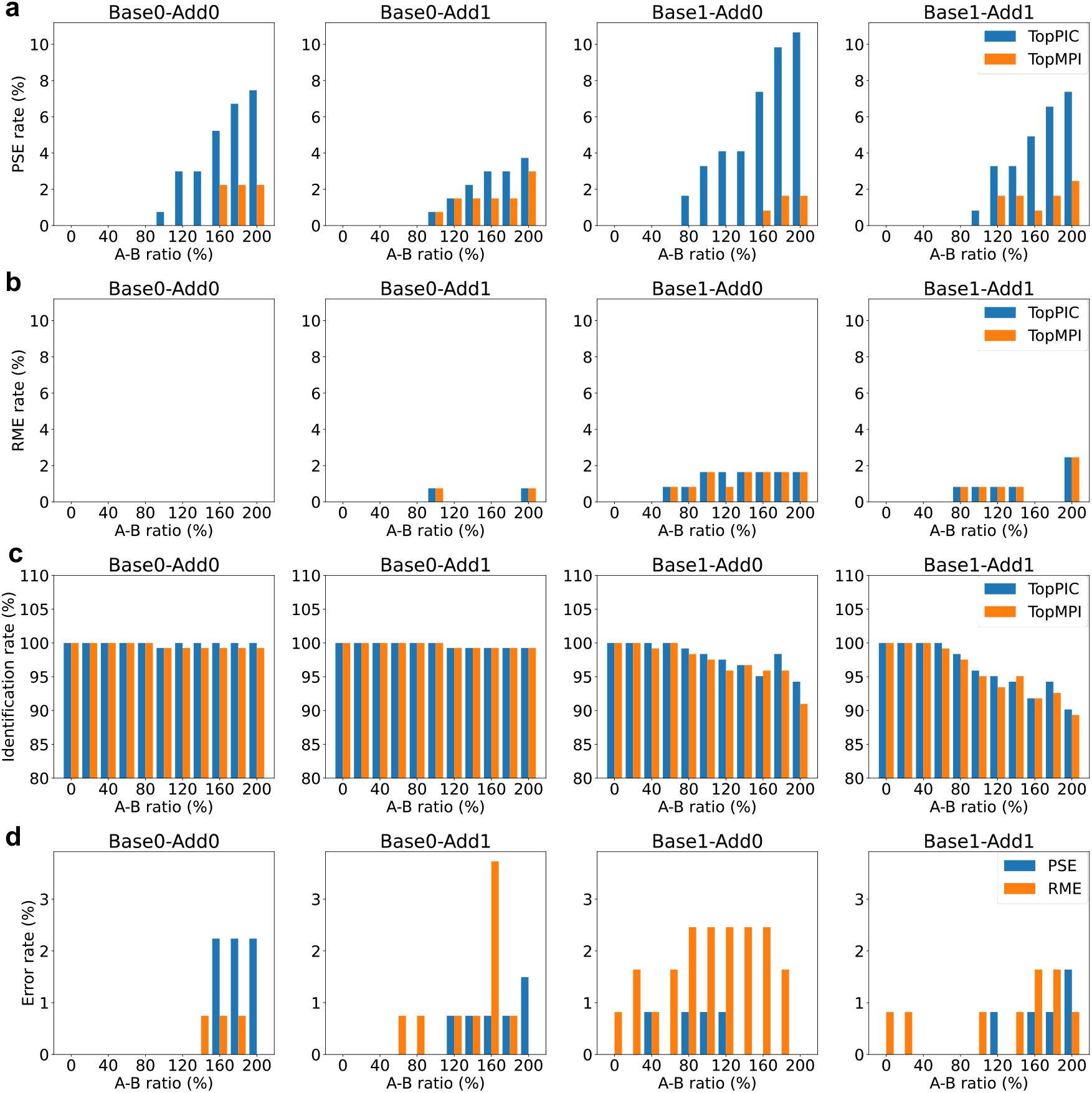
Error rates and identification rates of PrSMs reported by TopPIC and TopMPI at a 1% spectrum-level FDR across four types of pseudo-multiplexed spectra with varying A-B ratios (0%, 20%, …, 200%). (a) PSEs reported for base precursors. (b) RMEs reported for base precursors. (c) Identification rates for base precursors. (d) PSEs and RMEs reported for additive precursors by TopMPI.

We also created an evaluation dataset of pseudo-multiplexed MS/MS spectra by combining two non-multiplexed MS/MS spectra identified from the *E. coli* data set. A pair of spectra from the 498 non-multiplexed spectra was considered a matched pair if it met three criteria: (1) the charge states of their top SCPFs were different, (2) the difference between the average *m*/*z* values of their SCPFs was no more than 1.5 *m/z*, and (3) their proteoform identifications were from two different proteins. A total of 725 matched spectrum pairs were identified, and a pseudo-multiplexed MS/MS spectrum was generated for each pair by combining the fragment masses of the two spectra. The spectrum with a higher number of matched fragment masses was designated as the base spectrum with the most abundant precursor in the multiplexed spectrum, while the other spectrum was designated as the additive spectrum.

The 725 pseudo-multiplexed spectra were searched against the UniProt E. coli K12 proteome database (version September 7, 2023; 4,530 entries) concatenated with a shuffled decoy database of equal size using TopPIC and TopMPI (parameter settings provided in Supplemental Tables S2 and S5). PrSMs reported by the two tools were filtered with a 1% spectrum-level FDR. TopPIC identified 724 PrSMs, of which 2 (0.3%) were incorrect. TopMPI reported PrSM pairs for 713 spectra and single PrSMs for 12 spectra. Among the 1,438 single PrSMs reported by TopMPI, 1,432 (99.5%) were correct, 2 (0.2%) were PSEs, and 4 (0.3%) were RMEs. Notably, all PSEs were associated with base precursors, while all RMEs were associated with additive precursors. Further analysis of the E-value distribution of the original PrSMs used to construct the pseudo-multiplexed spectra revealed that TopMPI tends to miss PrSMs with low-confidence identifications in the original non-multiplexed spectra (Fig. 4).

**Fig. 4:**
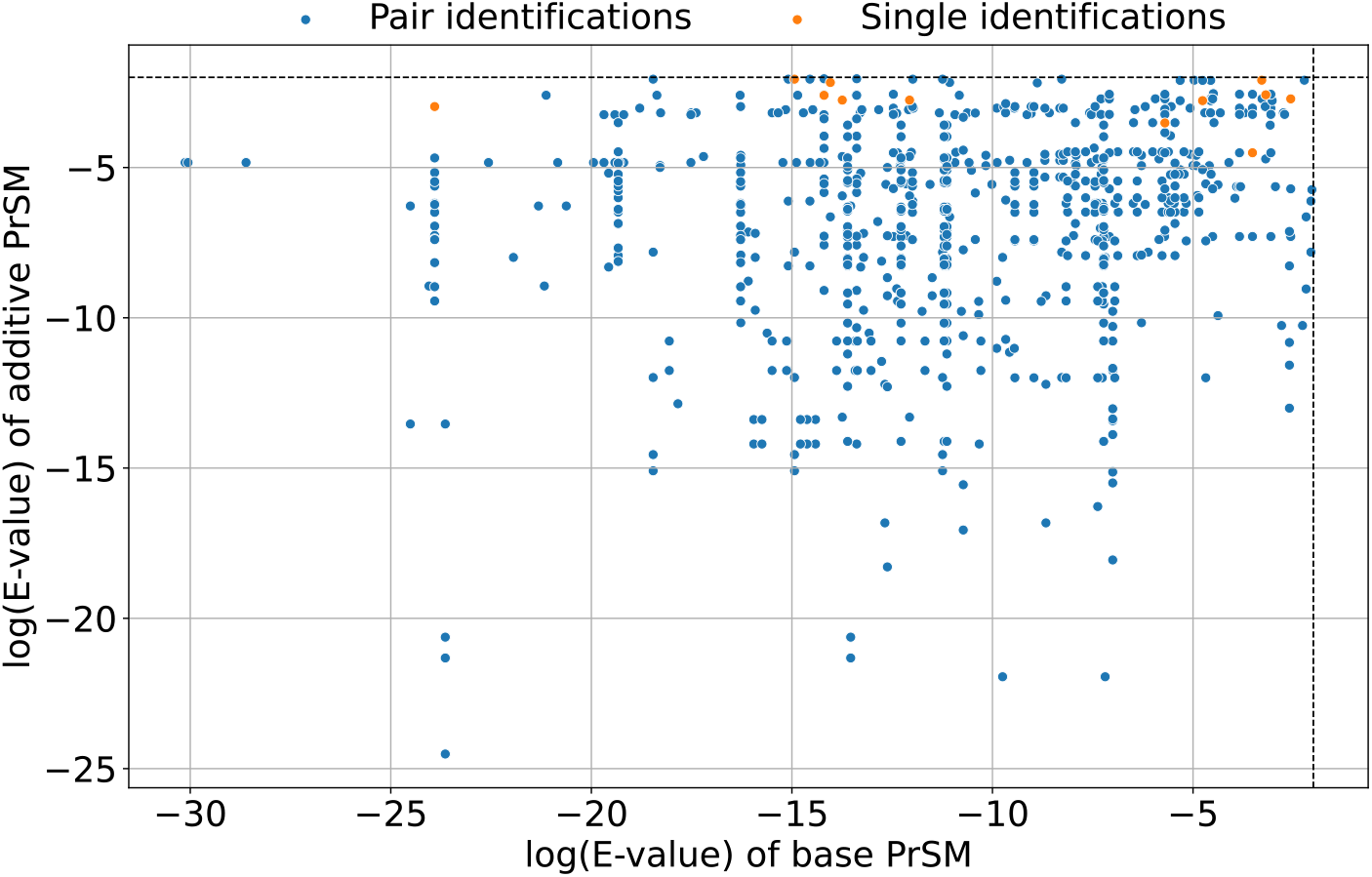
Distributions of log E-values (base 10) for the PrSMs of the base and additive non-multiplexed spectra in the 725 pseudo-multiplexed spectra. The E-values of the two PrSMs in each pseudo-multiplexed spectrum were reported by TopPIC. TopMPI identified at least one proteoform from each pseudo-multiplexed spectrum. Proteoform pairs identified by TopMPI are shown in blue, while single proteoform identifications are shown in orange.

### Comparison of TopMPI and TopPIC

We compared spectral identifications reported by TopMPI and TopPIC on the first and third replicates of a CZE-MS/MS data set of yeast proteins (Pride ID: PXD046651) [19]. The second replicate was excluded due to its low data quality. In the experiments, an LPA-coated capillary (50 μm i.d., 360 μm o.d., 1 m in length) was used for separation. MS/MS data were acquired using a Q-Exactive HF mass spectrometer in the data-dependent acquisition (DDA) mode. The MS and MS/MS spectra were collected at resolutions of 120,000 and 60,000 (at 200 *m*/*z*), respectively. The eight most intense precursor ions in each MS spectrum were isolated with a 2 *m*/*z* window to acquire higher-energy collision dissociation (HCD) MS/MS spectra. The first and third replicates contained 7,632 and 9,480 MS/MS spectra, respectively.

Following data preprocessing, TopPIC (version 1.7.6) and TopMPI were used to search the MS/MS spectra against the UniProt yeast proteome database (version March 3, 2023; 6,727 entries) with parameter settings detailed in Supplemental Tables S2 and S5. In both methods, PrSM and proteoform identifications were filtered using a 1% FDR. For replicate 1, TopPIC identified 4,544 PrSMs and 2,052 proteoforms, while TopMPI increased PrSM identifications by 25.9% (5,721 PrSMs) and proteoform identifications by 15.7% (2,374 proteoforms) (Fig. 5a). Among the TopMPI identifications, 4,622 PrSMs and 2,082 proteoforms were from primary precursors, 1,099 PrSMs and 675 proteoforms were from secondary precursors, and 383 proteoforms were shared by primary and secondary precursor identifications. Similarly, for replicate 3, TopPIC identified 4,422 PrSMs and 2,068 proteoforms, while TopMPI increased PrSM identifications by 26.3% (5,584 PrSMs) and proteoform identifications by 16.3% (2,406 proteoforms) (Fig. 5b). Of the TopMPI identifications, 4,489 PrSMs and 2,101 proteoforms were from primary precursors, 1,095 PrSMs and 662 proteoforms were from secondary precursors, and 357 proteoforms were shared by primary and secondary precursor identifications. Additionally, TopMPI demonstrated comparable reproducibility of proteoform identifications across the two replicates compared to TopPIC, with reproducibility rates of 62.0% versus 61.6%, respectively (Fig. 5c, 5d).

**Fig. 5:**
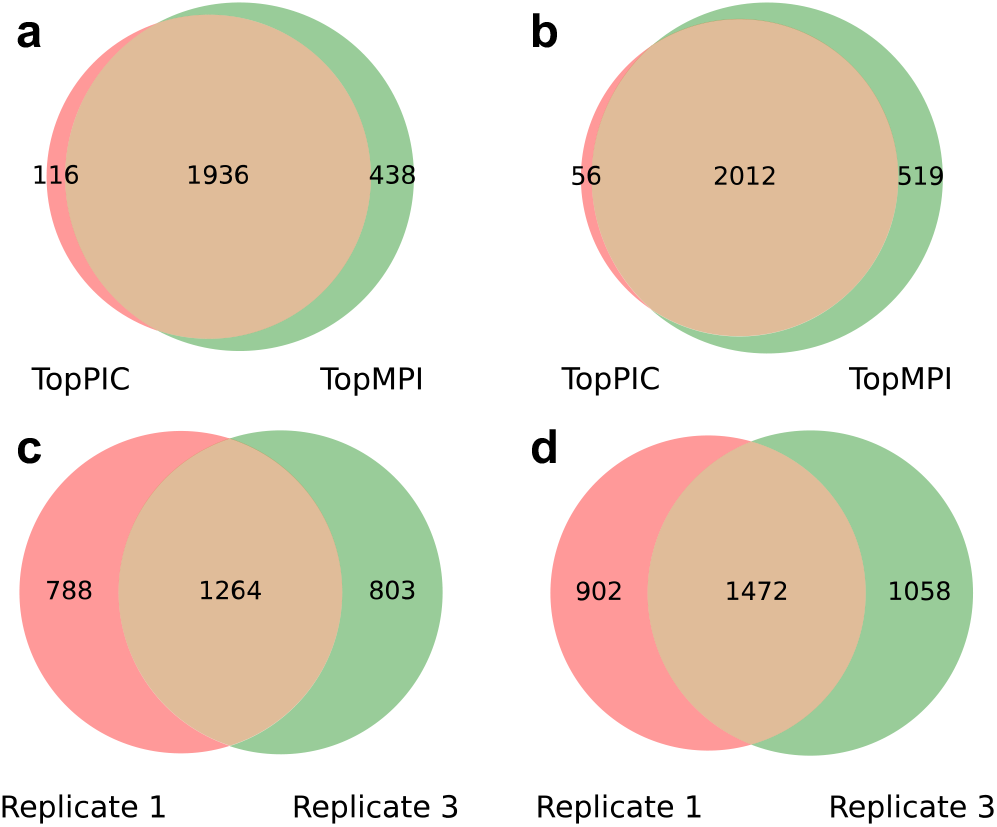
Comparison of proteoform identifications reported by TopPIC and TopMPI from replicates 1 and 3 of the yeast data set. Venn diagrams of proteoform identifications reported by TopPIC and TopMPI from replicate 1 (a) and replicate 3 (b). Venn diagrams of proteoform identifications reported from replicates 1 and 3 by TopPIC (c) and TopMPI (d). The ratios between the overlapping identifications and those reported from replicate 1 are 61.6% and 62.0% for TopPIC and TopMPI, respectively.

## Discussion and Conclusions

We developed TopMPI, a software tool for identifying multiplexed top-down mass spectra. TopMPI comprises two main functions. The first is determining the primary precursor within the isolation window, which is expected to generate the most fragment ions in the MS/MS spectrum. The second function employs a greedy approach, first identifying a PrSM using the primary precursor, then removing the matched fragment masses, and subsequently using the secondary precursor and the remaining fragment masses to identify another PrSM.

TopMPI reduces PSEs in PrSMs reported from multiplexed MS/MS spectra compared to existing tools designed for non-multiplexed spectra identification, such as TopPIC [15]. PSEs commonly arise when a multiplexed spectrum is incorrectly treated as a non-multiplexed spectrum during spectral identification. The precursor selection function in TopMPI effectively corrects many of the PSEs observed in TopPIC-reported identifications.

Beyond reducing PSEs, TopMPI also increases proteoform identifications compared to tools designed for non-multiplexed spectra identification. In top-down MS analysis of complex samples, it is common that multiple proteoforms are coeluted within the same isolation window. These multiplexed spectra are often treated as non-multiplexed, where only the precursor with the highest signal intensity is selected for spectral identification. This approach fails to identify proteoforms whose precursor intensities are not the highest within the isolation window. TopMPI enables the identification of these proteoforms, thereby increasing the sensitivity of proteoform identifications.

Despite its advantages, TopMPI has several limitations. First, it determines the primary precursor based solely on the number of NMFMs, which is insufficient to correct all PSEs. Incorporating chromatogram profile similarity between precursor and fragment masses could further reduce PSEs. Second, TopMPI employs a greedy approach for spectral identification, where the E-value of the primary precursor identification tends to be less significant due to the presence of fragment masses from the secondary precursor in the spectrum. This can lower the sensitivity of spectral identification. An alternative approach—directly matching a multiplexed spectrum to a combination of two proteoforms—may help address this issue and enhance spectral identification sensitivity.

## Code availability

The source code of TopMPI is available at https://github.com/todanielwang/TopMPI.

## Acknowledgements

This research was funded by NIH through the grants R01GM118470 and R01CA247863.

## Language polishing

The authors used ChatGPT to enhance the language and readability during the preparation of this paper. After utilizing ChatGPT, the authors reviewed and edited the content and take full responsibility for the final version of the paper.

## Competing interests

X.L. has a project contract with Bioinformatics Solutions Inc., a company that develops software for MS data processing.

