## Supplemental materials for "Proteoform identification using multiplexed top-down mass spectra"

Zhige Wang[1], Xingzhao Xiong[2], and Xiaowen Liu[2]

[1]Department of Computer Science, Tulane University, New Orleans, Louisiana, 70112, United States

[2]Deming Department of Medicine, Tulane University, New Orleans, Louisiana, 70112, United States

### Tables

**Table S1**: Parameter settings of TopFD

| Input Parameter | Value |
|---|---|
| Maximum charge | 30 |
| Maximum mass | 70,000 Da |
| m/z error tolerance of spectral peaks | 0.02 m/z |
| MS1 signal-to-noise ratio | 3 |
| MS/MS signal-to-noise ratio | 1 |
| Use MSDeconv score | False |
| ECScore cutoff | 0.5 |
| Number of MS1 scans to detect a feature | 3 |
| Use noise levels in single scans to filter | False |
| Disable final filtering of envelopes | False |
| Disable additional feature search | False |

**Table S2**: Parameter settings of TopPIC for "Precursor intensity and spectral identification"

| Input Parameter | Value |
|---|---|
| Fixed modification | Carbamidomethylation on cysteine |
| Allowed N-terminal forms | None, NME, NME_ACETYLATION, M_ACETYLATION |
| Maximum number of mass shift | 1 |
| Minimum value of a mass shift | -500 Da |
| Maximum value of a mass shift | 500 Da |
| Use a shuffled decoy database | True |
| Mass error tolerance | 10 ppm |
| Proteoform error tolerance | 1.2 Da |
| Spectrum level cutoff type | FDR |
| Spectrum level cutoff value | 0.01 |
| Proteoform level cutoff type | FDR |
| Proteoform level cutoff value | 0.01 |
| Use TopFD Features | True |

**Table S3**: Parameter settings of TopPIC for “Evaluation on pseudo-multiplexed MS/MS spectra”

| Input Parameter | Value |
|---|---|
| Fixed modification | Carbamidomethylation on cysteine |
| Allowed N-terminal forms | None, NME, NME_ACETYLATION, M_ACETYLATION |
| Maximum number of mass shift | 1 |
| Minimum value of a mass shift | -500 Da |
| Maximum value of a mass shift | 500 Da |
| Use a shuffled decoy database | False |
| Mass error tolerance | 10 ppm |
| Proteoform error tolerance | 1.2 Da |
| Spectrum level cutoff type | E-value |
| Spectrum level cutoff value | 0.01 |
| Proteoform level cutoff type | E-value |
| Proteoform level cutoff value | 0.01 |
| Use TopFD Features | True |

**Table S4**: Parameter settings of TopMPI for evaluating $\delta$ and $\gamma$

| Input Parameter | Value |
|---|---|
| Fixed modification | Carbamidomethylation on cysteine |
| Allowed N-terminal forms | None, NME, NME_ACETYLATION, M_ACETYLATION |
| Maximum number of mass shift | 1 |
| Minimum value of a mass shift | -500 Da |
| Maximum value of a mass shift | 500 Da |
| Use a shuffled decoy database | True |
| Mass error tolerance | 10 ppm |
| Proteoform error tolerance | 1.2 Da |
| TopPIC spectrum level cutoff type | E-value |
| TopPIC spectrum level cutoff value | 10,000 |
| TopMPI spectrum level cutoff type | FDR |
| TopMPI spectrum level cutoff value | 0.01 |
| TopMPI use TopFD Features | Yes |
| TopMPI proteoform level cutoff type | FDR |
| TopMPI spectrum level cutoff type | 0.01 |
| $\alpha$ | 0.2 |
| $\beta$ | 0.9 |
| $\gamma$ | Various settings |
| $\delta$ | Various settings |

**Table S5**: Parameter settings of TopMPI for comparing with TopPIC

| Input Parameter | Value |
|---|---|
| Fixed modification | Carbamidomethylation on cysteine |
| Allowed N-terminal forms | None, NME, NME_ACETYLATION, M_ACETYLATION |
| Maximum number of mass shift | 1 |
| Minimum value of a mass shift | -500 Da |
| Maximum value of a mass shift | 500 Da |
| Use a shuffled decoy database | True |
| Mass error tolerance | 10 ppm |
| Proteoform error tolerance | 1.2 Da |
| TopPIC spectrum level cutoff type | E-value |
| TopPIC spectrum level cutoff value | 10,000 |
| TopMPI spectrum level cutoff type | FDR |
| TopMPI spectrum level cutoff value | 0.01 |
| TopMPI use TopFD Features | Yes |
| TopMPI proteoform level cutoff type | FDR |
| TopMPI spectrum level cutoff type | 0.01 |
| $\alpha$ | 0.2 |
| $\beta$ | 0.9 |
| $\gamma$ | 4 |
| $\delta$ | 5 |

**Figure**

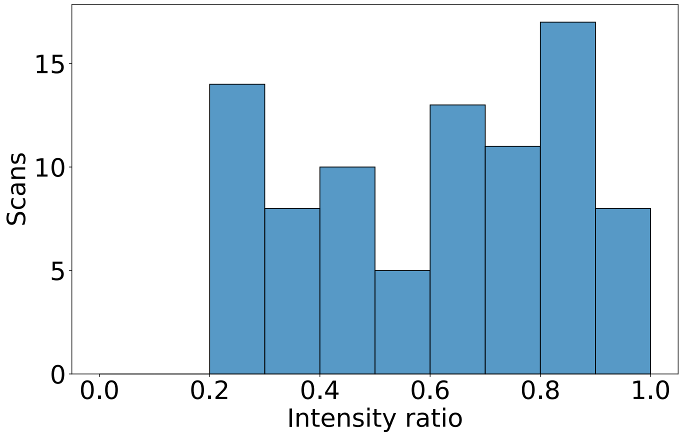


**Fig. S1**: The distribution of the intensity ratio of the second and first most abundant precursors in the 86 MS/MS spectra with proteoform pair identifications reported from the *E. coli* data set. The minimum value of the intensity ratio is 0.21.
